# Development and parameterisation of a size-structured multispecies bioeconomic model integrating consumer demand

**DOI:** 10.64898/2026.09.04.749338

**Authors:** Emy Cottrant, Nicolas Barrier, Bruno Ernande, Sophie Gourguet, Alaia Morell, Hanna Schenk, Yunne-Jai Shin, Martin Quaas

## Abstract

A high proportion of the world’s population relies on marine fisheries as a source of food and employment, highlighting the need for sustainable exploitation strategies. However, fisheries management commonly relies on single-species models that overlook ecological interactions and economic trade-offs, which may lead to stocks being exploited above sustainable levels. To address this gap, we developed a novel size-structured, multispecies bioeconomic module integrated within the spatially explicit, individual-based OSMOSE ecosystem model. OSMOSE represents exploited fish communities in which individuals interact through opportunistic, size-dependent predator–prey interactions which are explicitly incorporated into the bioeconomic framework. The bioeconomic model accounts for multiple species and size classes, allowing for size-dependent market prices. Fishing costs and profits are represented using an extension of the Gordon-Schaefer model to multiple interacting species, while consumer demand is modelled using a nested Dixit-Stiglitz utility function with three levels of constant elasticity of substitution for fish commodity, species and size classes. We present a method for estimating parameters for the bioeconomic module when empirical estimates are unavailable, using the North Sea OSMOSE configuration, which comprises 15 species. Our method successfully estimated the six bioeconomic parameters required to operationalise the model. Results indicate that differences in cost parameters were primarily associated with variability in species biomass rather than fishing gear, while fish prices were more strongly influenced by consumer’s demand than by availability. This framework provides a basis for assessing the economic consequences of alternative climate change scenarios and supporting sustainable fisheries management.

## 1. Introduction

Fisheries and marine harvesting support billions of people worldwide through employment or food supply (OECD 2025). The consumption of aquatic food worldwide has increased at a rate of 3% per year on average since 1960, while the human population has grown by 1.6% which highlights the increasing use of fish products (FAO 2024). Climate change and over-fishing are threatening the stability of many marine and freshwater ecosystems (Allison et al. 2009; Sumaila et al. 2011; Cheung & Sumaila 2015), with less than two thirds of fish stocks exploited at sustainable levels in 2021 (FAO 2024), thus, raising concerns about food security and the future economic income derived from fisheries. Three pillars of sustainability have been identified by the United Nations, namely economic development, social development and environmental protection, which should be considered in management systems (Asche et al. 2018). Approaches to sustainable fisheries management have been gradually combining biological and economic components into single models in order to maintain food security and ecosystems (Nielsen et al. 2018). One widely employed framework is the Gordon-Schaefer (G-S) model for commercial fisheries (Gordon 1954; Schaefer 1957). Based on a simple logistic representation of biomass dynamics, this model allows deriving the maximum sustainable yield, along with the maximum economic yield, as it also accounts for harvesting costs and market prices. However, the G-S model assumes constant prices, thus ignoring the effect of changing supply on the market prices of fish. A more realistic approach is to specify a downward-sloping demand function (Dao et al. 2023), which allows prices to respond to variations in supply and better reflects economic behavior, while also including broader social objectives required by the United Nations for sustainable fisheries. In a multi-species framework, the substitutability between different species of fish further plays a role in market dynamics (Lus & Muriel 2009). One approach to take this into account in fisheries bio-economic modeling is to specify a utility function of the Dixit and Stiglitz (1977) type, which assumes a constant elasticity of substitution between different fish species (Quaas & Requate 2013).

While bioeconomic models are increasingly used, multiple limitations remain, mainly in data availability which prevent the global use of those models. In fact, economic data availability for fisheries is highly variable depending on the area of investigation. For example, in the European Union, Annual Economic Reports (AER) provide all necessary data such as landings, fishing costs and revenues (STECF 2024), but only for a small, and not necessarily representative, subset of fleets. Also, in some regions, such data are not collected or not publicly accessible (Lam et al. 2011). In addition, many studies focus on a limited number of species, typically those highly impacted by fisheries, often ignoring key interactions such as trophic, technical and economic ones. For example, the Atlantic cod *Gadus morhua*, has been widely studied in isolation to improve the management of its fisheries in the North-East Atlantic, despite being embedded in complex food webs (Reeves et al., 2014) and frequently exploited in mixed fisheries (Schenk et al. 2023; Han et al. 2025). Furthermore, while some models have included multiple species, catches are often aggregated instead of being partitioned into size or age classes, thereby overlooking size-dependence that significantly influences both ecological interactions (Barnes et al. 2010) and market prices (Zimmermann et al. 2011; Voss et al. 2022). Accounting for these different types of interactions is important in order to provide more accurate population estimates that will be used, not only to set appropriate fishing quotas, but also for improving fisheries management while ensuring sustainable revenues for fishers.

Here, we present an adapted version of the bioeconomic G-S model that accounts for fishing revenues, exploitation costs, and a consumer demand system that incorporates multiple species and size classes of fish. This framework allows fish prices to vary as a function of supply and consumers’ preferences for species and size. The model is designed to be flexible enough to adapt to data availability, representing different types of fisheries. For some species, size-structured data may be lacking, while for others, data are available at the commercial category level, reflecting different market values depending on fish size. We describe the equations of the model and present an application to the multi-species ecosystem model OSMOSE (Shin and Cury 2004). The goal is to investigate parameter estimation in a multispecies bio-economic model, with a particular focus on how consumers’ preferences across species and size classes shape fish prices.

## 2. Method

### 2.1. Model description

We developed and applied a multispecies size-structured bioeconomic model accounting for costs of fishing along with consumer’s demand. The model requires estimates of harvested biomass and accessible biomass (*i.e.* biomass composed of individuals larger than the minimum size harvested) for each species considered in the model, with biomass being size-structured according to commercial sales categories. This allows the model to be coupled with any size-based multispecies model possessing the required output. The OSMOSE model (Shin and Cury 2004) is used here as an application, but the model can use other sources, such as annual fisheries stock assessments (e.g. stock assessment reports from the International Council for the Exploration of the Sea; ICES 2020).

First, we are describing equations accounting for costs and profits of fishing for each species *i* and each size-class *s* over time *t*. Fishing cost *C* depends on the stock accessible biomass *B* and the harvested biomass *H*, with costs increasing with harvested biomass, and decreasing when accessible biomass increases. Thus, cost of fishing is represented by

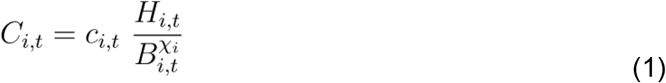

with *H*_*i,t*_ and *B*_*i,t*_ representing the sum of harvested biomass *h*_*i,s,t*_ and accessible biomass *b*_*i,s,t*_ over size classes *s* following 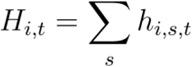 and,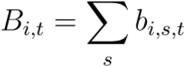 respectively. χ_*i*_ ≥ 0 is the stock elasticity of species *i* where χ*i*<1 represents hyperstability, thus, if the fish stock decreases, fishing costs increase less than proportionally. 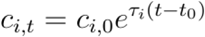 represents the baseline cost of each species over time where *c*_*i*_,0 is the baseline cost at the beginning when *t*=*t*_0_ and *T*_*i*_ represents the temporal trend on fishing cost. The temporal trend parameter combines numerous causes of fishing price fluctuation such as trend on fish sell prices (Willmann and Kelleher 2010), external factors such as competition from aquaculture, but also variation of the fuel cost, and technical progress (improved fishing gear will harvest more efficiently).

Profit Π_*i,t*_ is defined here for each species *i* as the difference between revenue and fishing cost *C*_*i,t*_ (1), and represented by

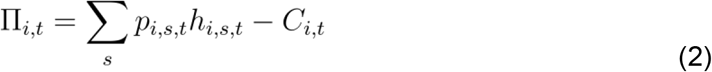

with *P*_*i,s,t*_ representing the price for size-class *s* of species *i* at time *t*

The profit margin π_*i,t*_ is the fraction of profits (2) with respect to revenues

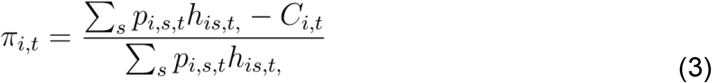

Consumer’s utility for fish consumption is described by a nested constant elasticity of substitution (CES) utility structure (Figure 1). At the highest level, consumer’s utility is derived from two inputs: consumption of fish or other food. Consumption of fish is then derived from the consumption of each species *i* available to the consumer. Finally, the consumption of a species *i* is derived from the consumption of each size class *s* available to the consumer for this species.

**Figure 1.**
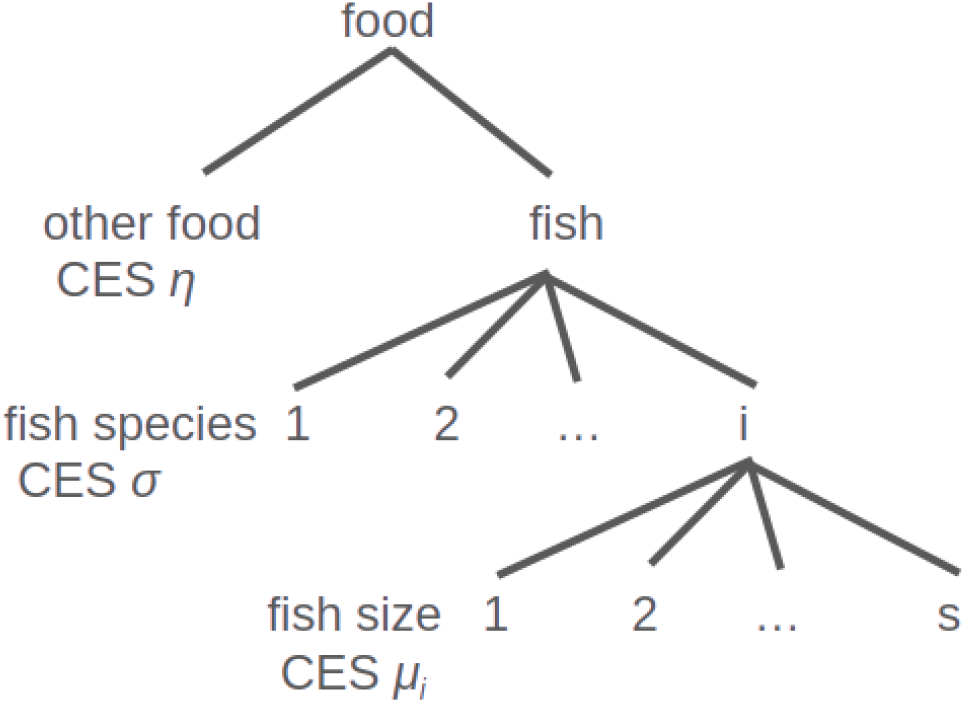
Nested CES structure of consumer’s demand for fish.

Consumers pay for fish, thus, the total utility of fish consumption *v*_*t*_ over another commodity is described by the total utility function where the cost of fish is subtracted to the total benefit of consuming fish.

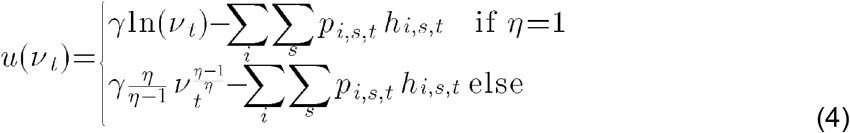

Where γ >0 represents the total expenditure on fish, η >0 is the elasticity of demand for fish compared to another commodity.

The utility of consumption of a fish species *v*_*t*_ is adapted from the Dixit-Stiglitz utility function (Dixit and Stiglitz 1977), where species are considered imperfect substitutes for each other, represented by a constant elasticity of substitution:

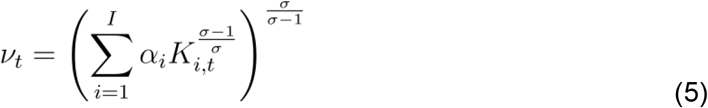

Where σ >0 represents the elasticity of substitution between different fish species *i*. A high value of σ can be associated with consumers’ flexibility in substituting one fish species for another, while a low σ represents a strong consumer’s preference for a stable distribution of fish species composition in the consumption basket (Quaas and Requate 2013). We assume σ > η since different species of fish are better substitutes than other commodities (Quaas and Requate 2013; Schenk et al. 2023). *αi*, represents the consumer’s relative preference for a species *i* with 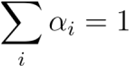 . The gain from fish consumption *v*_*t*_ also depends on the gain from the consumption of each size class of each species, described by the utility function *K*_*i,t*_ . Similarity to *v*_*t*_, the size-classes dependant utility *K*_*i,t*_ is derived from the Dixit-Stiglitz utility function (Dixit and Stiglitz 1977) where size classes are considered imperfect substitutes for each other for each species:

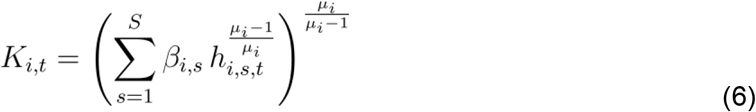

μ_*i*_ represents the elasticity of substitution between size-classes where μ_*i*_> σ as a consumer will first substitute for another size-class of the same species more easily than between species. β_*i,s*_ represents the consumer’s relative preference for a size-class *s* of species *i*, as bigger fish might have higher consumption value than small ones (Zimmermann and Heino 2013, Schenk et al. 2023), with 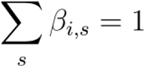 .

The inverse demand function is obtained by maximizing utility *u*(*V*_*t*_) with *v*_*t*_from equation (4) (See Quaas and Requate 2013 for details on the derivation of the inverse demand function), resulting in

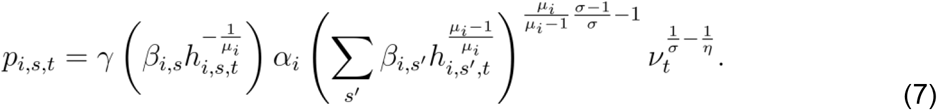

### 2.2. Parameter estimation

To estimate the parameters from the fishing cost equation, the method by Voss *et al.* (2022) is used. The profit margin (3) is rearranged with equation (1) as follows:

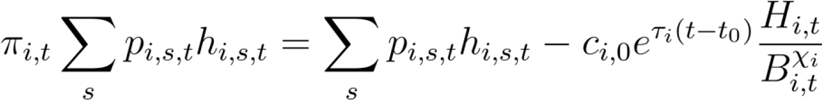

After log transforming, we further obtain:

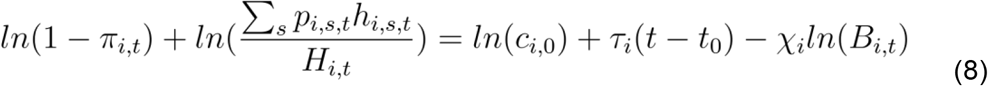

Parameters accounting for baseline cost *c*_*i*,0_, trend on fishing cost *T*_*i*_ and stock elasticity *X*_*i*_ are obtained by providing species selling prices *p*_*i,s,t*_, harvested biomass *h*_*i,s,t*_ and accessible biomass over time *b*_*i,s,t*_, profit margins over time π_*i,t*_ and by using a multiple linear regression.

To estimate *β*_*i,s*_ and *α*_*i*_ for each commercially size-structured species, we rearrange the inverse demand function (7) (Supplementary material A1) to obtain firstly for the estimation of *β*_*i,s*_

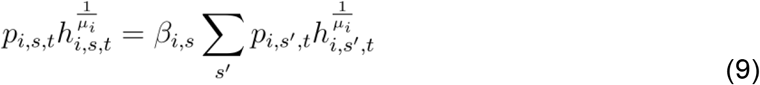

And secondly for the estimation of *α*_*i*_

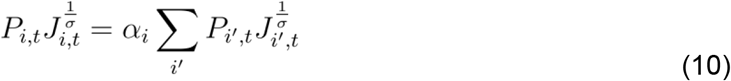

with 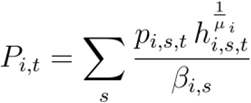 and 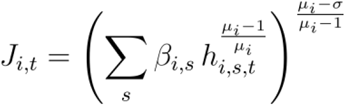

Furthermore, to estimate *γ* and *η* we rearranged the inverse demand function (8) to obtain

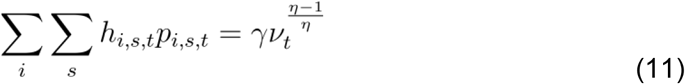

### 2.3. Case study

We applied the bioeconomic model previously stated to the North Sea region comprising ICES area 4 and 7d, the North Sea and the eastern English Channel, respectively. Based on total fisheries landings (ICES 4abc and 7d) and the scientific North Sea International bottom trawl survey (NS-IBTS-Q1, DATRAS), 15 species of interest were chosen, representing the major economically important teleosts species caught in the North Sea (Morell et al. 2023; Table 1). Species of interest were composed of seven demersal species, three benthic species and five pelagic species. To provide the best estimate possible for our bioeconomic model in order to better predict future income from fisheries, we needed a model that provides size-structured accessible biomass and landings for each species to match commercial sales categories, along with additional mortality due to trophic interaction and the life cycle of the species (e.g. larval mortality, starvation). Thus, we chose the OSMOSE model, a spatially explicit multi-species, size-structured and individual-based model, driven by trophic interactions, growth, reproduction and explicit mortality from predation, starvation and fishing (see Shin and Cury 2004 for details). Here we used the North Sea-Eastern English Channel configuration, using Bioen-OSMOSE, described and parameterized in Morell *et al.* (2023). Based on the original OSMOSE framework, Bioen-OSMOSE also includes the emergence of life history traits due to bioenergetic variations in abiotic and biotic factors (See Morell *et al.* 2023 for details).

**Table 1:**
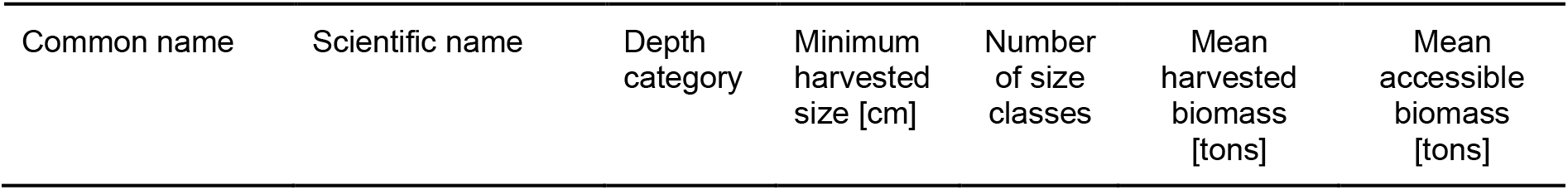
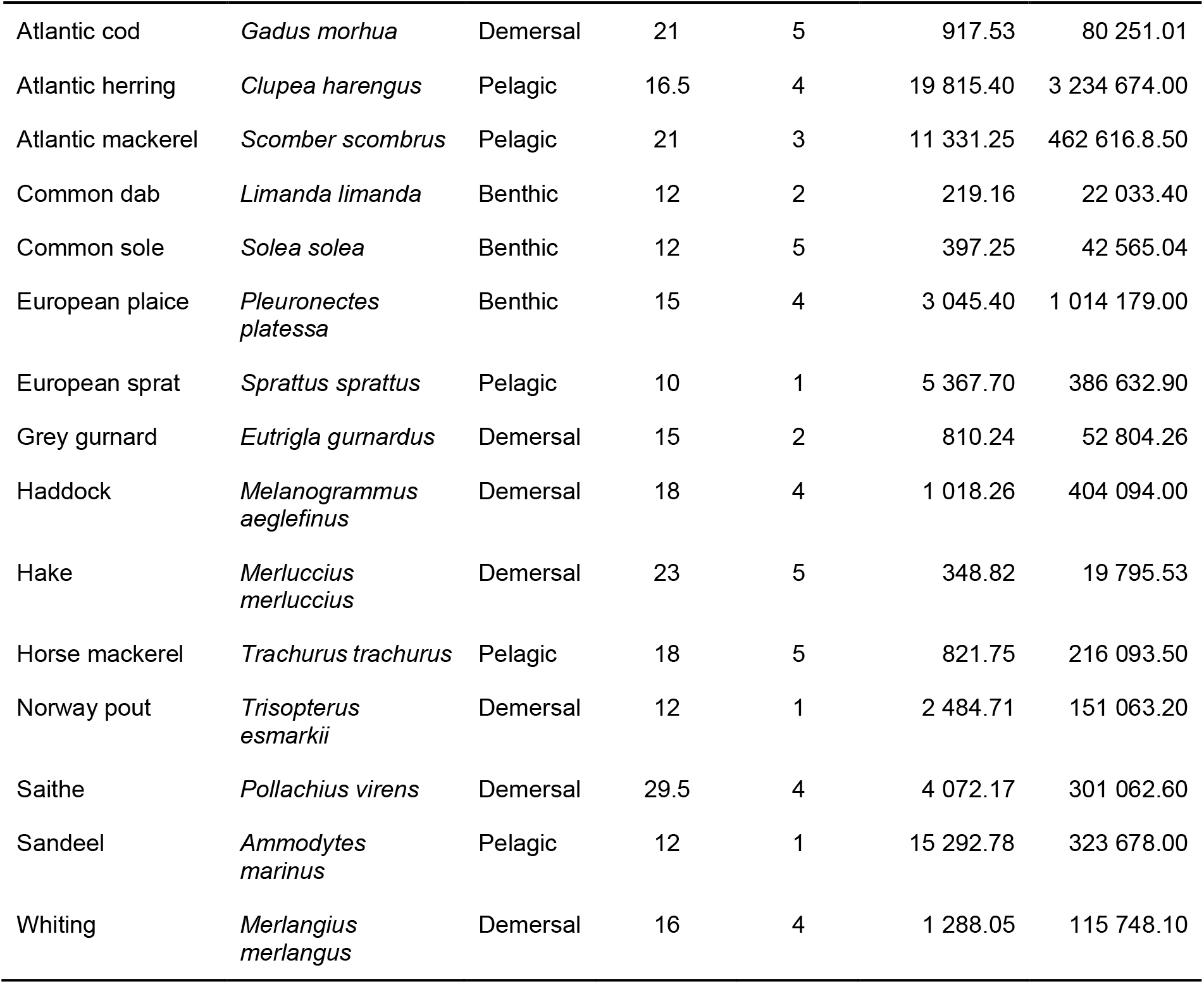
List of species included in the North Sea OSMOSE configuration with minimum harvested size (in cm), number of size classes used for the economic model, and mean harvested biomass and accessible biomass in tons for a year (2010-2019).

| Common name | Scientific name | Depth category | Minimum harvested size [cm] | Number of size classes | Mean harvested biomass [tons] | Mean accessible biomass [tons] |
| --- | --- | --- | --- | --- | --- | --- |
| Atlantic cod | *Gadus morhua* | Demersal | 21 | 5 | 917.53 | 80 251.01 |
| Atlantic herring | *Clupea harengus* | Pelagic | 16.5 | 4 | 19 815.40 | 3 234 674.00 |
| Atlantic mackerel | *Scomber scombrus* | Pelagic | 21 | 3 | 11 331.25 | 462 616.8.50 |
| Common dab | *Limanda limanda* | Benthic | 12 | 2 | 219.16 | 22 033.40 |
| Common sole | *Solea solea* | Benthic | 12 | 5 | 397.25 | 42 565.04 |
| European plaice | *Pleuronectes platessa* | Benthic | 15 | 4 | 3 045.40 | 1 014 179.00 |
| European sprat | *Sprattus sprattus* | Pelagic | 10 | 1 | 5 367.70 | 386 632.90 |
| Grey gurnard | *Eutrigla gurnardus* | Demersal | 15 | 2 | 810.24 | 52 804.26 |
| Haddock | *Melanogrammus aeglefinus* | Demersal | 18 | 4 | 1 018.26 | 404 094.00 |
| Hake | *Merluccius merluccius* | Demersal | 23 | 5 | 348.82 | 19 795.53 |
| Horse mackerel | *Trachurus trachurus* | Pelagic | 18 | 5 | 821.75 | 216 093.50 |
| Norway pout | *Trisopterus esmarkii* | Demersal | 12 | 1 | 2 484.71 | 151 063.20 |
| Saithe | *Pollachius virens* | Demersal | 29.5 | 4 | 4 072.17 | 301 062.60 |
| Sandeel | *Ammodytes marinus* | Pelagic | 12 | 1 | 15 292.78 | 323 678.00 |
| Whiting | *Merlangius merlangus* | Demersal | 16 | 4 | 1 288.05 | 115 748.10 |

The biological unit of the model is a school, representing individuals from the same species, born simultaneously, and biologically identical (e.g. age, location, weight, abundance). The time step of OSMOSE is two weeks. Harvested biomass *h*_*i,s,t*_ and accessible biomass *b*_*i,s,t*_, expressed in tons, are directly extracted from OSMOSE with the model being calibrated according to 2010-2019 fisheries landings, size-at-age from scientific surveys and estimated biomass for assessed species (Morell et al. 2023). Accessible biomass is defined as the total biomass of individuals whose total length is above the minimum size allowed in catches. Multiple OSMOSE processes are stochastic such as species movement and mortality (see Shin and Cury 2004 for details), therefore, 20 replicates of the model were run, and parameters were estimated for each run resulting in 20 replicates for each parameter.

To estimate the bioeconomic cost parameters, species prices *p*_*i,s,t*_ were extracted from the EUMOFA database for the 2010-2019 period with size classes based on the commercial categories described in the common marketing standards of the European Council Regulation (1996) (Table 2). Prices corresponded to monthly first sale prices of full fresh fish, and prices of gutted fresh fish were also added if not enough data was available. Thus, in the following analysis, consumers refer to buyers at fish markets but retail prices paid by end consumers are expected to be correlated to first sale prices, as each intermediary typically applies a constant percentage markup. In the North Sea OSMOSE configuration, fisheries are not explicitly represented, thus, fishing mortality is applied per species. Nevertheless, for each species, fishing mortality takes into account size dependent selectivity. Profit margins *π*_*i*_ were obtained for the same time period, with one profit margin per year, based on the Scientific, Technical and Economic Committee for Fisheries AER data report (STECF 2024). Due to the discrepancy between STECF AER report, presenting landings per species per fleet for each ICES area, profit per fleet for the North Atlantic region (NAO), and OSMOSE output presenting harvested biomass per species, the profit margin was obtained by firstly rescaling the costs of fishing to obtain the profit associated to the North Sea region (ICES 27.4 and 27.7d; hereafter named NS) for each fleet and then by calculating the weighted mean of profit per species. In detail, the profit margin comprised the gross value of landing per species to which was deducted the energy cost, the other variable costs and personnel costs associated with each species (Table 2). As energy costs and variable costs depends on the fishing effort while personnel costs depends on the value of landings, rescaling the costs associated with each fleet to the NS region was calculated separately using

**Table 2:** Sources of the variables needed for the parameter estimation associated with the North Sea OSMOSE configuration. NAO represents the North Atlantic Ocean and NS represents the North Sea region including ICES zones 27.4 and 27.7d.

| Variable | Description | Type of data | Source |
| --- | --- | --- | --- |
| $p_{i,s,t}$ | Price of species $i$ in euro/kg per size-class $s$ over time $t$ | Monthly | EUMOFA first sale prices |
| $V_{i,f,NS},$<br>$V_{i,f,NAO}$ | Gross value of landing for species $i$ of fleet $f$ in NS and NAO region, respectively, in euro. | Annual | STECF AER economic and transversal data |
| $Ce_{f,NAO}$ | Energy cost for fleet $f$ in NAO region, in euro. | Annual | STECF AER economic and transversal data |
| $Co_{f,NAO}$ | Other variable costs for fleet $f$ in NAO region, in euro. | Annual | STECF AER economic and transversal data |
| $Cp_{f,NAO}$ | Personnel costs for fleet $f$ in NAO region, in euro. | Annual | STECF AER economic and transversal data |
| $L_{i,f,NS}$ | Landings in kg for species $i$ of fleet $f$ in NS region | Annual | STECF AER economic and transversal data |
| $E_{f,NS},$<br>$E_{f,NAO}$ | Effort of fleet $f$ in days at sea for the NS and NAO regions, respectively. | Annual | STECF AER economic and transversal data |

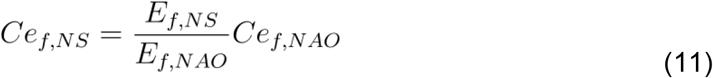

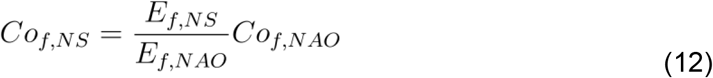

with *Ce*_*f,NAO*_ and *Ce*_*f,NAO*_ representing the energy costs and other variable costs, respectively, for each fleet *f* in the NAO region. *E*_*f,NS*_ and *E*_*f,NAO*_ representing the effort of each fleet *f* in the NS and NAO regions, respectively.

For the personnel costs associated to each fleet *f*, value of landing was used following

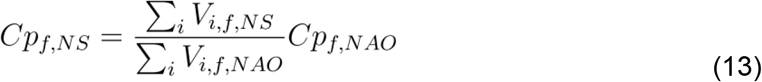

with *Cp*_*f,NAO*_ representing the personnel cost of each fleet *f* in the NAO region. *V*_*i,f,NAS*_ and *V*_*i,f,NAO*_ representing the gross value of landing of the fleet *f* for the species in the NS and NAO regions respectively.

Now, we have costs associated with each fleet in the NS region and we need to rescale them to obtain costs associated with each species within those fleets. Thus, for energy costs and other variable costs, we use the proportion of landing corresponding to the species of interest following

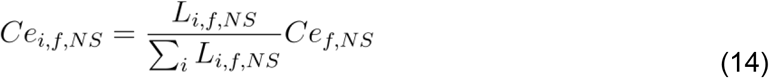

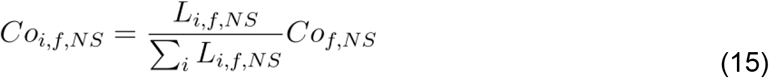

with *Ce*_*f,NAS*_ and *Co*_*f,NAS*_ from (11) and (12), respectively. *L*_*i,f,NS*_ represents landings in kg for each fleet *f* in the NS region.

For personnel costs, the same method was used but the gross value of landing was used instead of the weight of landing following

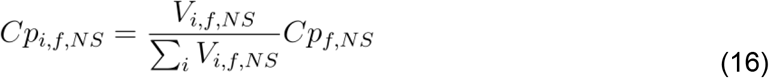

with *Cp*_*f,NAS*_ (13) and *V*_*i,f,NS*_ representing the gross value of landing of species *i* of fleet *f* in the NS region.

Then, all costs were summed over fleet *f* and profit margin was obtained following the formula

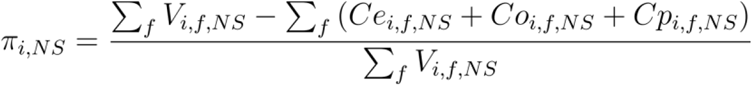

The method was applied for each year of the period 2010-2019 obtaining ten values of profit *π* for each species over time.

Estimations of *c*_*i*,0_, *X*_*i*_ and *T*_*i*_ were conducted by solving (8) for each replicate using a linear regression with the ‘lm’ function in R (R Core Team 2024). As the stock elasticity *X*_*i*_ needs to be between 0 and 3, the value was fixed to zero if the parameter estimation resulted in a negative value and fixed to values from Harley et al. (2001) if the value estimated was greater than 3. Because baseline cost *c*_*i,0*_ does not have a unit a reference cost *c*_*i,0,ref*_ was calculated using 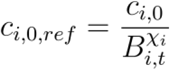 giving a cost for each species in euros. Three Generalized Linear Mixed Models (GLMMs) were applied, using the *lme4* package (Bates et al. 2015), to investigate the influence of species on estimated values of *c*_*i,0,ref*_, *X*_*i*_ and *T*_*i*_ with the replicate number as a random effect. After analysis, species were grouped according to depth categories (i.e. benthic, demersal and pelagic) which could help link values to different fishing techniques such as pelagic trawlers, purse seine or bottom trawlers.

For the reference cost *c*_*i,0,ref*_, values were strictly positive, thus, a gamma family was used with an identity link following the formula:

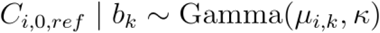

where *µ*_*i,k*_ = E(*C*_*i,0,ref*_| *b*_*k*_) and *k >* 0 is the shape parameter. The conditional variance was

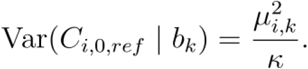

Using an identity link, the mean response was modelled as

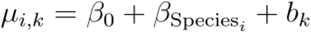

where *β*_0_is the intercept (corresponding to the reference level of Species), 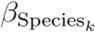 represents the fixed effect associated with species *i*, and *b*_*k*_ is a random intercept associated with the replicate *k*.

Random effects were assumed to follow a normal distribution:

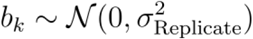

For the trend on fishing cost, the GLMM used a gaussian family with identity link following the formula:

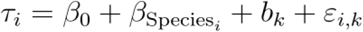

where *β*_0_is the intercept corresponding to the reference level of Species, 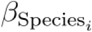 represents the fixed effect of Species *i*, and *b*_*k*_ is a random intercept associated with replicate *k*.

Random effects and residual errors were assumed to follow normal distributions:

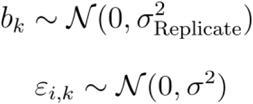

For the stock elasticity *X*_*i*_, the GLMM used a gamma family with a log link following the formula:

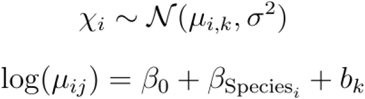

where β_0_is the intercept corresponding to the reference level of Species, 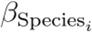 represents the fixed effect of Species *i*, and *b*_*k*_ is a random intercept associated with replicate *k* following:

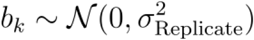

Estimations of β_*i,s,*_ andα_*i*_ were obtained by solving (9) and (10), respectively, because only one parameter was being estimated for each equation, we used a linear regression with the “lm” function from R (R Core Team 2024). Values were then compared with proportions of harvested biomass and fish prices considering that higher prices would be found for preferred species and size class. For β_*i,s,*_, proportions of harvested biomass and fish prices were calculated as the harvested biomass and prices per size class, respectively, divided by the sum of harvested biomass and prices over size classes. For α_*i*_, the method was similar for the proportion of harvested biomass, where the harvested biomass for a species was divided by the sum over species. For the proportion of fish prices, firstly, for each size class the price was ponderated by the catches with the price of a size class being multiplied by the proportion of harvested biomass of that size class. Then, the prices were summed over size class and the proportion of price was obtained by dividing the price of a species by the sum over species of fish prices.

Finally, γ was obtained by solving (11) using a linear regression with the “lm” function from R (R Core Team 2024). For all estimations, parameters η,σ µ_*i*_ and were fixed to 4, 6 and 8 respectively (Schenk et al. 2023), following the condition µ_*i*_>σ>η previously stated.

## 3. Results

Due to the stochasticity of the OSMOSE model, parameter estimations for the cost function (8) had high differences between replicates for some species, resulting in high mean and SE, especially for baseline cost *c*_*i,0*_, thus the median is used for*c*_*i*,0_,> while the mean is used for the other parameters (Table 3). Baseline cost *c*_*i,0*_, ranged between 132.91 and 2.43×10^21^ with a median across species and replicate of 6 149.49 with a mean of 8.09×10^18^ (Table 3). The species with the lowest median baseline cost over 20 replicates was European sprat *Sprattus sprattus* with 132.91 while the species with the highest median baseline cost was the European plaice *Pleuronectes platessa* with 8.93×10^6^ (Table 3). After rescaling *c*_*i,0*_, to obtain the reference cost *c*_*i,0*_,_*ref*_, the cost ranged from 105.32 and 4 405.41 euro.ton^-1^ with a mean of 1 192.85 euro.ton^-1^ ± 58.65 SE (Table 3). The species with the lowest mean *c*_*i,0*_,_*ref*_, over 20 replicates was European sprat with 131.98 euro.ton^-1^ while the species with the highest mean baseline cost was the common sole *Solea solea* with 4 319.70 euro.ton^-1^ (Table 3). The benthic category exhibited the highest mean *c*_*i,0*_,_*ref*_, with 2 402.49 euro.ton^-1^ ± 310.16 SE, followed by demersal species with 1 271.96 euro.ton^-1^ ± 107.50 SE and pelagic species exhibited the lowest *c*_*i,0,ref*_, with 356.31 euro.ton^-1^ ± 35.63 SE. Significant differences were found between some species but globally, looking at depth categories, differences were showing outliers within the model but no clear difference was found between groups (Figure 1A).

**Figure 1.**
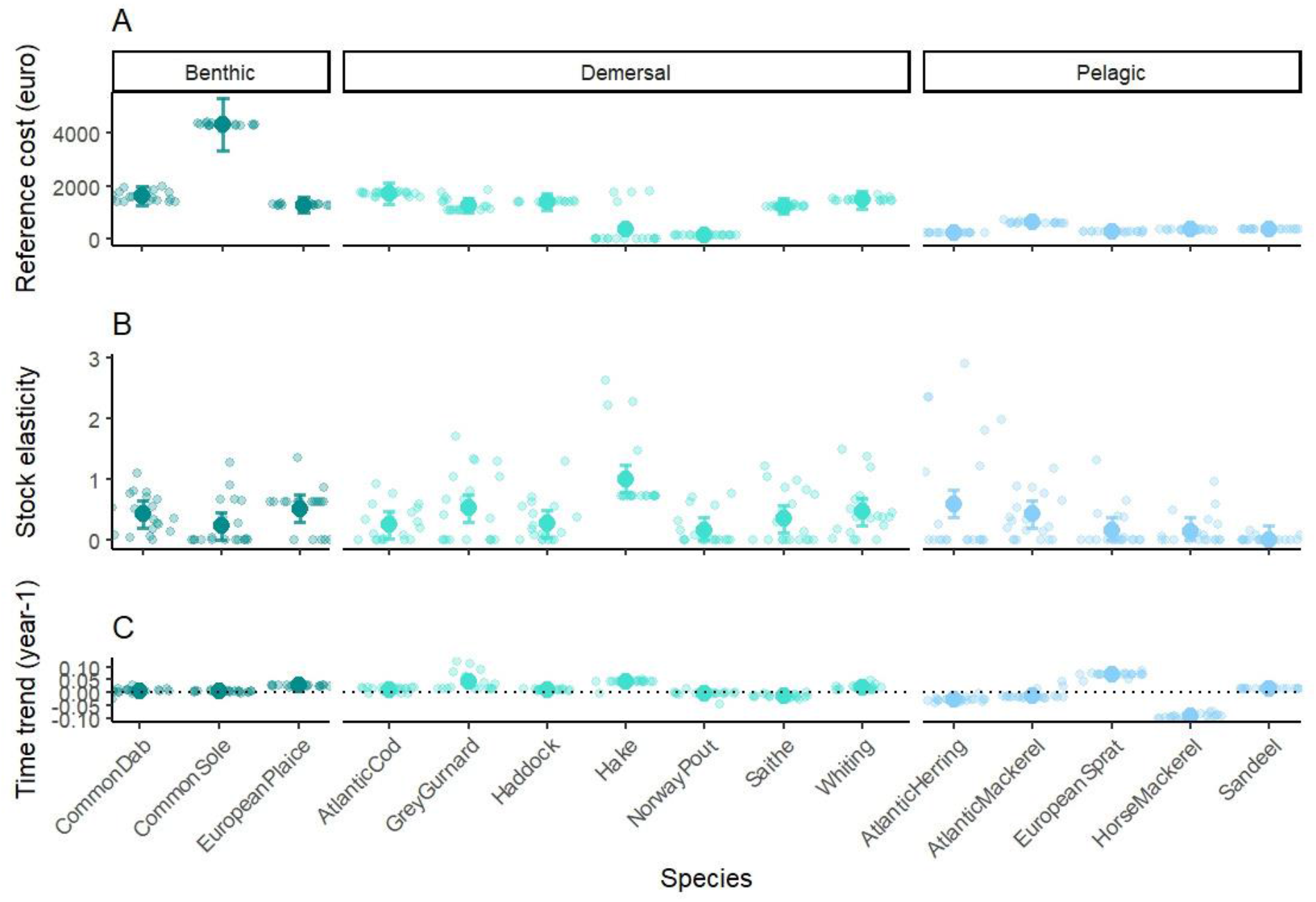
Average reference cost (A), stock elasticity (B) and time trend (C) estimated by the GLMM for each species and depth category with error bars representing standard deviation, and points representing raw data.

**Table 3:** Estimated parameters values for cost functions. Values presented represent the median of 20 replicates for baseline cost *c*_*i,0*_ with the mean in brackets. And the mean of 20 replicates for the reference cost *c*_*i,0,ref*_, stock elasticity *X* and trend on fishing cost *τ*_*i*_, with the SE in brackets. * represents species that had *τ*_*i*_ changing signs across replicates.

| Species name | $C_{i,0}$ | $C_{i,0,ref}$ | $\chi_i$ | $\tau_i$ (year<sup>-1</sup>) |
| --- | --- | --- | --- | --- |
| Atlantic cod | 11 020.05 (3.58x10<sup>6</sup>) | 1 706.44 (15.97) | 0.263 (0.064) | 0.012 (0.002)* |
| Atlantic herring | 222.10 (1.21x10<sup>20</sup>) | 222.62 (1.77) | 0.600 (0.214) | -0.028 (0.002) |
| Atlantic mackerel | 22 958.22 (9.45x10<sup>12</sup>) | 612.28 (10.69) | 0.432 (0.111) | -0.011 (0.004)* |
| Common dab | 68 829.03 (6.51x10<sup>6</sup>) | 1 605.09 (41.72) | 0.432 (0.067) | 0.007 (0.002)* |
| Common sole | 4 397.71 (1.75x10<sup>8</sup>) | 4 319.70 (12.17) | 0.246 (0.083) | 0.004 (0.001)* |
| European plaice | 8.93x10<sup>6</sup> (8.82x10<sup>9</sup>) | 1 282.68 (5.53) | 0.527 (0.077) | 0.027 (0.001) |
| European sprat | 273.96 (2.61x10<sup>8</sup>) | 271.06 (2.79) | 0.161 (0.073) | 0.071 (0.002) |
| Grey gurnard | 55 798.38 (2.77x10<sup>10</sup>) | 1 263.97 (51.67) | 0.534 (0.127) | 0.041 (0.008) |
| Haddock | 10 568.54 (1.26x10<sup>9</sup>) | 1 400.63 (4.70) | 0.279 (0.074) | 0.011 (0.001) |
| Hake | 2.35x10<sup>5</sup> (7.73x10<sup>12</sup>) | 1 688.99 (99.79) | 1.015 (0.134) | 0.038 (0.002) |
| Horse mackerel | 338.43 (3.21x10<sup>6</sup>) | 336.06 (2.34) | 0.155 (0.057) | -0.094 (0.002) |
| Norway pout | 132.91 (9.06x10<sup>4</sup>) | 131.98 (1.48) | 0.162 (0.059) | -0.003 (0.002)* |
| Saithe | 4 598.42 (3.91x10<sup>8</sup>) | 1 230.99 (7.60) | 0.361 (0.095) | -0.014 (0.001) |
| Sandeel | 338.71 (521.30) | 339.54 (0.79) | 0.016 (0.009) | 0.014 (0.001) |
| Whiting | 4.36x10<sup>5</sup> (4.36x10<sup>9</sup>) | 1 480.73 (13.74) | 0.479 (0.096) | 0.019 (0.002) |

Estimations for the stock elasticity *X*_*i*_ showed high differences between species ranging from 0.000 to 2.911 (mean= 0.378 ± 0.029 SE; Table 3). The species that exhibited the lowest mean *X*_*i*_ over 20 replicates was sandeel *Ammodytes marinus* with 0.016; while the species that exhibited the highest mean *X*_*i*_ was hake *Merluccius merluccius* with 1.015 (Table 3). Looking at differences between groups of species, the demersal category had the highest mean *X*_*i*_ with 0.442 ± 0.043, followed by benthic species with 0.402 ± 0.047, and pelagic species had the lowest mean *X*_*i*_ with 0.273 ± 0.056 (Figure 1B). Results showed that differences between some species were found, no significant difference between depth categories was identified.

Taking a look at the trend *τ*_*i*_ in fishing price values, seven species exhibited slightly positive values for all the replicates, showing that (nominal) cost of fishing increased over time. Three species, Atlantic herring *Clupea harengus,* horse mackerel *Trachurus trachurus* and saithe *Pollachius virens*, had negative values for all replicates, and five species exhibited both negative and positive values across replicates (Table 3). Globally, the trend on fishing price ranged between -0.106 and 0.124 year^-1^ (mean= 0.006 year^-1^ ± 0.002 SE; Table 3). The species that exhibited the lowest mean *τ*_*i*_ was horse mackerel *Trachurus trachurus* with -0.094 year^-1^, while the species that exhibited the highest *τ*_*i*_ was European sprat with 0.071 year^-1^. When comparing depth categories, the demersal category exhibited the highest *τ*_*i*_ with a mean of 0.015 year^-1^ ± 0.002 SE, followed by the benthic category with 0.012 year^-1^ ± 0.002 SE, and pelagic species exhibited the lowest *τ*_*i*_ with a mean of -0.010 year^-1^ ± 0.005 SE. Results showed that differences between depth categories were explained by species variability, especially within the pelagic group where both the minimum and the maximum were *τ*_*i*_ found (Figure 1C).

Results for consumer’s size preference *β*_*i,s*_ were differing depending on the species. Consumer’s preferences *β*_*i,s*_ were not always higher for larger size classes as only five species had their highest size preference matching their highest size (41.7%), and four species had their highest size preference corresponding to their second to highest size (33.3%; Figure 2). It can be observed that, as expected, the price *p*_*i,s,t*_ is higher for higher size preference (Figure 2). A difference in trend between *β*_*i,s*_ and *p*_*i,s,t*_can be observed only for the Atlantic mackerel *Scomber scombrus*, where, while *p*_*i,s,t*_ was increasing for the highest size, *β*_*i,s*_ decreased due to a strong decrease in harvested biomass *h*_*i,s,t*_ (Figure 2).

**Figure 2.**
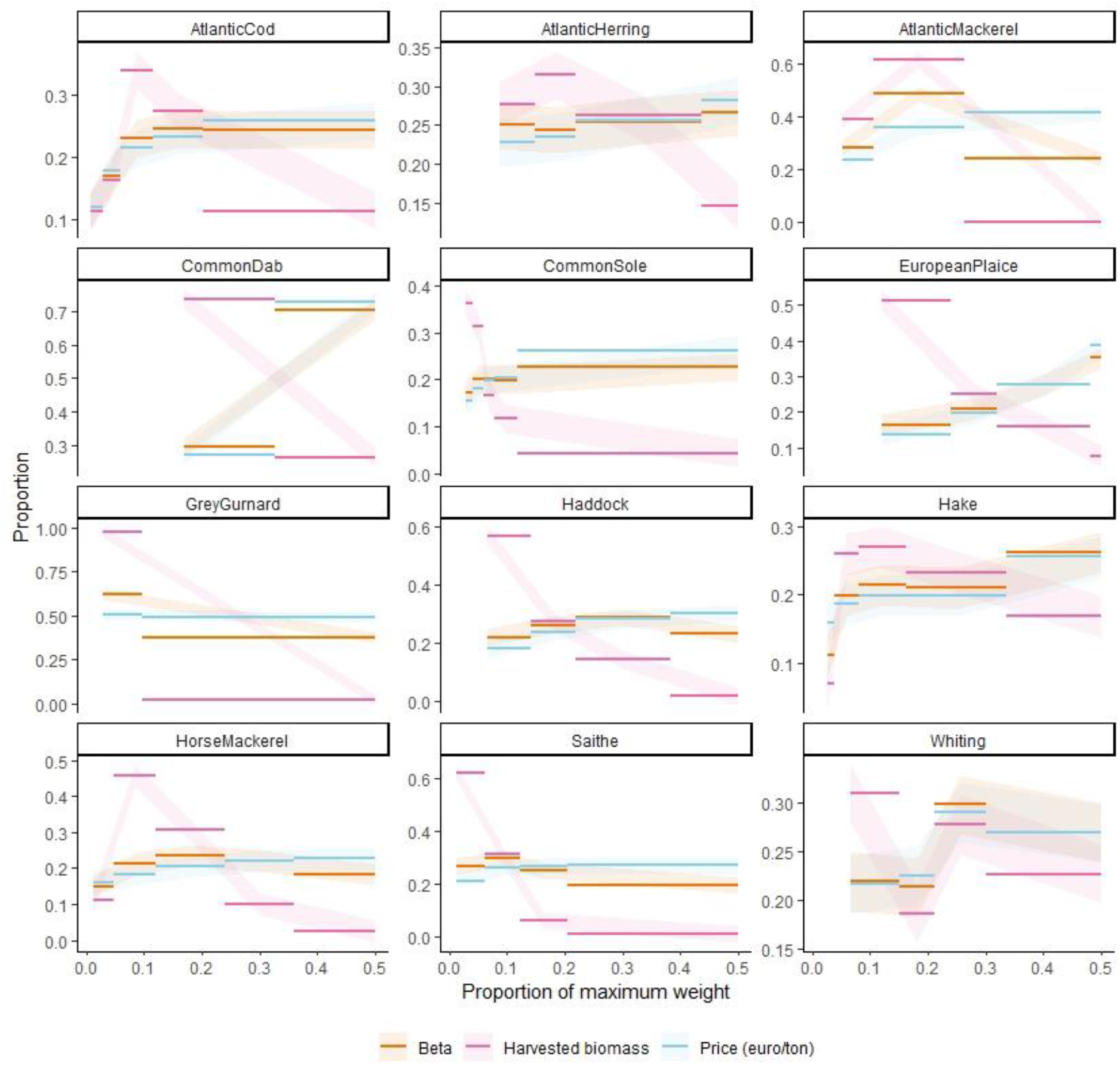
Mean estimation of consumer’s size preference *β*_*i,s*_ (orange) and mean harvested biomass *h*_*i,s,t*_ (pink) over 20 replicates along with price *p*_*i,s,t*_ (blue) for each commercial category. Commercial categories were represented as the proportion of the maximum weight of each species. As *β*_*i,s*_ is a proportion between 0 and 1, harvested biomass and prices were represented as proportions of the total harvested biomass and the total selling price of species, respectively. As variability between replicates was extremely low (*ca.* 0.001), ribbons were set to 0.02 and used only to indicate the direction of data for comparison.

Species preference *α*_*i*_ showed a consumer’s preference for Common sole followed by Atlantic cod, hake and haddock *Melanogrammus aeglefinus*, while lower preference was found for Norway pout *Trisopterus esmarkii*, European sprat and Grey gurnard *Eutrigla gurnardus* (Figure 3). We compared the estimated value for *α*_*i*_ to the proportion of harvested biomass and proportion of fish prices and observed that *α*_*i*_ was closer to the proportion of fish prices and that, regardless of a high increase in harvested biomass, the value of *α*_*i*_ remained low for species such as Atlantic herring and Sandeel.

**Figure 3.**
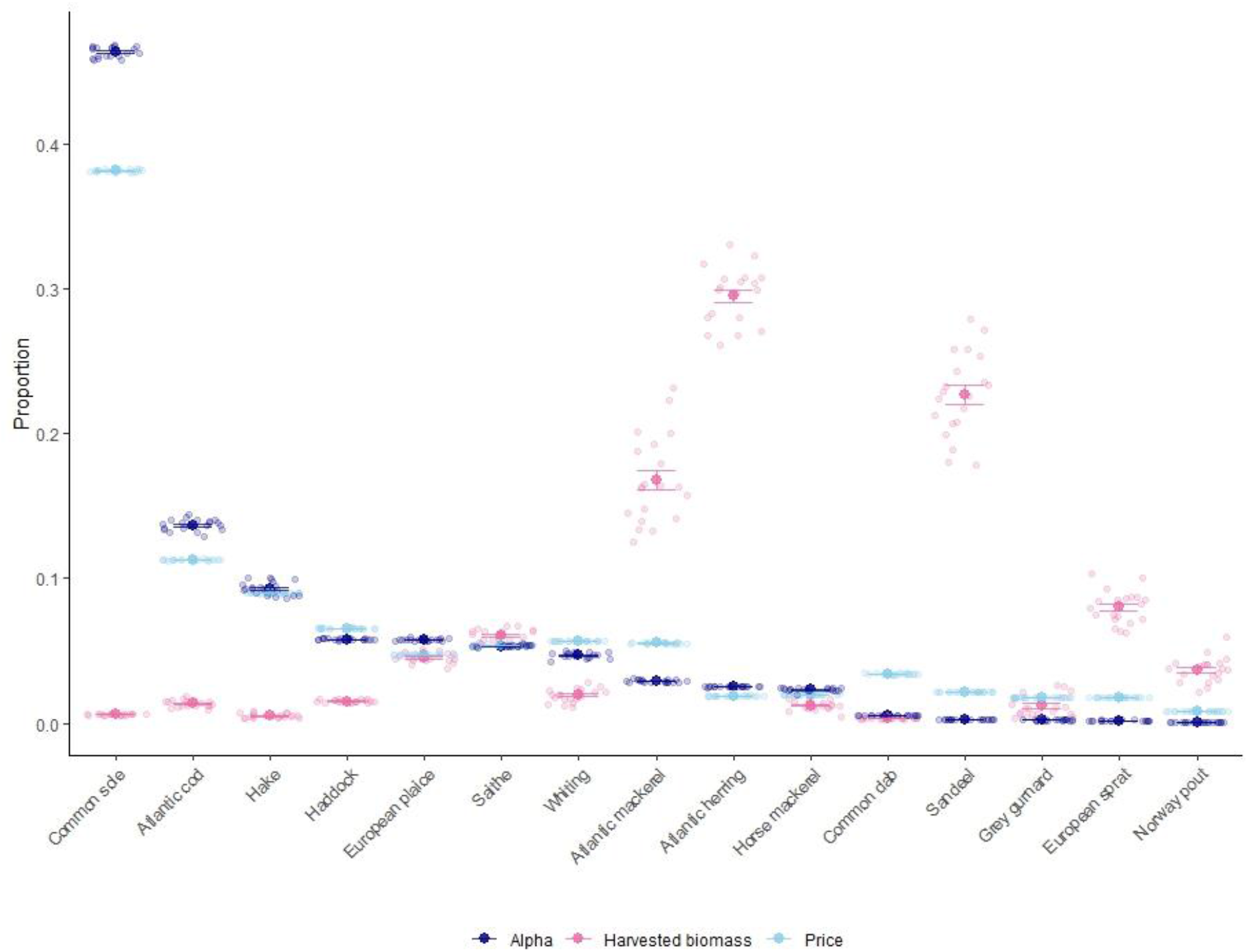
Mean over replicate of the consumer’s species preference *α*_*i*_ (in red) with grey lines representing each of the 16 replicates.

The total expenditure on fish γ ranged between 687 407 and 746 224 with a mean of 718 958.3 ± 3 321.6 SE.

Globally, results of the parameter estimation showed that the demand system assumes that consumers’ preferences increases the price associated with that size class or species.

## 4. Discussion

Our study presents a method for the estimation of bioeconomic parameters for modeling future costs of fishing, profits, social welfare and fish prices. A total of six different parameters were estimated per species and per replicate resulting in 1 800 estimations. Using biomass, catch and fish size outputs from the OSMOSE model applied to the North Sea, we successfully obtained estimation for economic parameters that cannot be found in the literature, which is particularly true for less commercially important species such as Norway pout and Grey gurnard. In fact, profits of fisheries are proven to be extremely difficult to estimate in most parts of the world as a lot of fishermen are reluctant to share the different costs associated with their activities (Daurès et al. 2013), or data are sometimes not made publicly available (Lam et al. 2011). Some exceptions exist such as in the European Union where details of fishing costs and fishing profits are available per fleet segment along with catches per species (STECF 2024), but at an aggregated level. Therefore, our model can be adapted to any ecosystem or fishing fleet but requires, at minimum, market prices data. In that case expert guesses for profit margin may substitute actual data.

For all fishing cost parameters (i.e. baseline cost, stock elasticity and trend on fishing cost), some differences were found between species but globally, there was no clear grouping factor between species explaining those differences. It was expected that differences in parameters could be observed between depth categories as species from the same depth category could be harvested by the same fishing gear. However, our results show no difference between depth groups which can be explained by the fact that the OSMOSE model is species specific and each species can be harvested by multiple engines regardless of their depth category. Regarding stock elasticity, our results corresponded to previous study by Harley et al. (2001) using ICES reports in multiple areas across the North Atlantic region, including the North Sea, for three species (*i.e.* European plaice, Whiting *Merlangius merlangus* and Saithe), while results were different for Atlantic cod, Haddock, Hake and Common sole. This could be due to the change in species accessible biomass between the time series used in both studies and, for the case of Hake, the value for gadiformes was used multiple times during the parameter estimation as the value estimated was higher than 3, thus, it lowered the mean result for this species. Almost all our species, except Hake, exhibited stock elasticity lower than one, which suggest hyperstability, a result often found when comparing catch-per-unit-effort (CPUE) to abundance (Peterman and Steer 1981; Crecco and Overholtz 1990; Hilborn and Walters 1992). While hyperstability is advantageous for fishermen because it ensures continuous catches regardless of a decrease in stock size, it is far from sustainable as a stock can collapse without any warning signal resulting from a decrease in harvested biomass (Hilborn and Walters 1993). For example, this was previously observed for the Peruvian anchoveta *Engraulis ringens* and Atlantic herring where collapse was not detected through a decrease in capture rate (Hilborn and Walters 1993). Previous literature also associated low values of stock elasticity close to 0 to pelagic species, usually presenting a schooling behavior (Ulltang 1980; Butterworth 1981; Bjørndal 1987), and higher values closer to 1 to demersal or benthic species as they often exhibit a more sparse distribution (Schaefer 1957), but this pattern was not found in our results. This could be due to stock elasticity being highly dependent on the stock data used which could explain the differences in parameter estimation for some species. For example, the stock elasticity of Atlantic cod was lower than that reported by Voss et al. (2022) and Ekerhovd and Gordon (2013) who used data from the Baltic Sea between 1974 and 2020 and from the North Atlantic between 1977 and 2011, respectively.

Our result indicates a majority of positive trends in fishing costs showing that over time, costs increase which could induce a decrease of profit for fisheries. This can be associated with an increase of price of fuel as seen in recent years, which not only affects how much a fishing trip costs but also affects the cost of fishing gears as most of their production relies on fuel (Willmann and Kelleher 2010). Regardless of prices of fuel, nominal costs can also increase due to inflation, and/or due to increasing wages. All those increases in costs are balanced by improved technologies (Squires and Vestergaard 2013), which help find fish schools with more efficiency, and more selective gears that catch relatively more target individuals and reduce bycatch. In our case study, we also found species that exhibited a negative trend in fishing costs (Atlantic herring, Horse mackerel and Saithe), this could be due to fluctuations in selling prices of those species over time, or increasing profit margin due to advances in technology that may reduce costs such as labor, maintenance and repair, and fuel.

Our parameter estimation showed a consumer’s preference for larger sizes. Only a limited amount of studies have investigated consumers’ preferences for fish size classes (Zimmermann et al. 2011; Zimmermann and Heino 2013) and, so far the relationship between price and size class have been investigated without estimating the full demand system. However, for some species, we found differing results showing that the biggest fish is not necessarily preferred. For example, we found a non linear relationship between preference and size class for Atlantic mackerel with the preferred size category being smaller than the maximum size. In fact, Atlantic mackerel is most commonly commercialized in smaller sizes with the medium size (between 200 and 500g) often preferred as it ensure a better plate coverage and a better resistance to heat during cooking (Mayol 2025). In addition, consumer’s preference could be constrained by availability of large fish as, due to fishery induced mortality towards larger sizes, it is possible that over time, abundance of larger size classes had decreased drastically, as observed for the Atlantic cod (Han et al. 2025).

Looking at species preference, results showed that some species are clearly preferred. In a previous study investigating how fish are sold and priced, Gallegati et al. (2011) found that harvested biomass globally did not influence the selling prices as buyers are not aware of the quantity available for sale. Thus, prices for the same species tend to go down within a day as the buyer realises that the quantity available for sale is sufficient, also known as the “declining price paradox”. While the harvested biomass did not directly influence consumer’s preference for a species, it is expected that over time, a high preference for a species could influence its sustainability. Including this parameter in future multi-species models would help with sustainable management, for example by implementing eco-labelling or educating consumers about the status of fish stocks, that would redirect consumptions towards more sustainable species and fishing practices (Menozzi et al. 2020; Kersulec et al. 2024). Moreover, our application showed global species preferences in the North Sea but it would be expected that if data was separated per country, preferences would differ as consumers from different countries rely on different criteria while choosing a fish to consume (Menozzi et al. 2020).

Our study presented equations of a multi-species size-structured bioeconomic model along with a case study using an OSMOSE model applied to the North Sea. We successfully estimated six bioeconomic parameters for each of the 15 modelled species. We also showed that multispecies bioeconomic models could be coupled to complex end-to-end models, taking into account predator-prey interactions and the influence of abiotic factors, aiming to support management decisions. Using this new modelling framework, future studies could for example estimate maximum economic yields from a multispecies perspective or explore climate change impacts on the dynamics of size-structured interacting populations and bioeconomic outcomes.

## Acknowledgements

This work was funded by France Filière Pêche through the ADAPT project (grant agreement PH/2022/10).

## Supplementary material

A1: Proof of equation 9 and 10

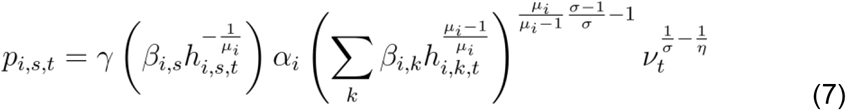

Multiplying by 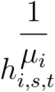,

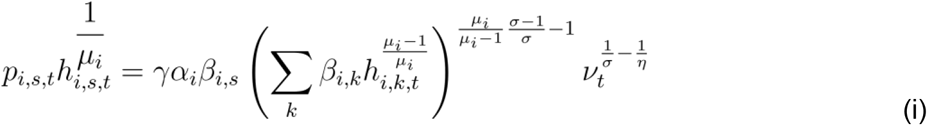

Summing over size classes,

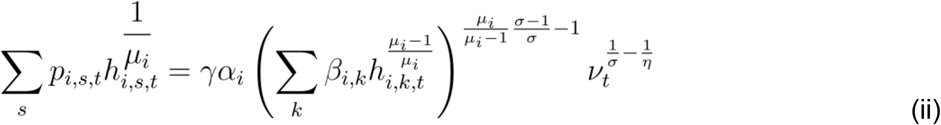

as 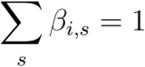 and using (ii) in (i),

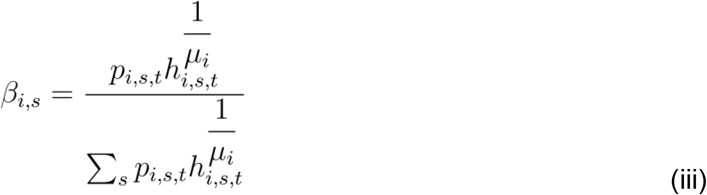

Multiplying (ii) by 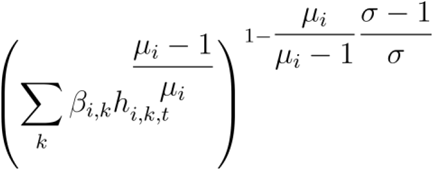,

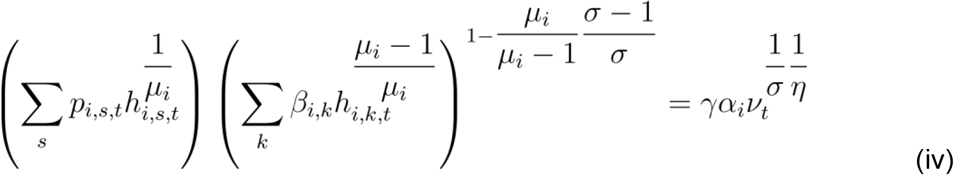

Summing over species and using 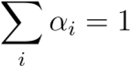,

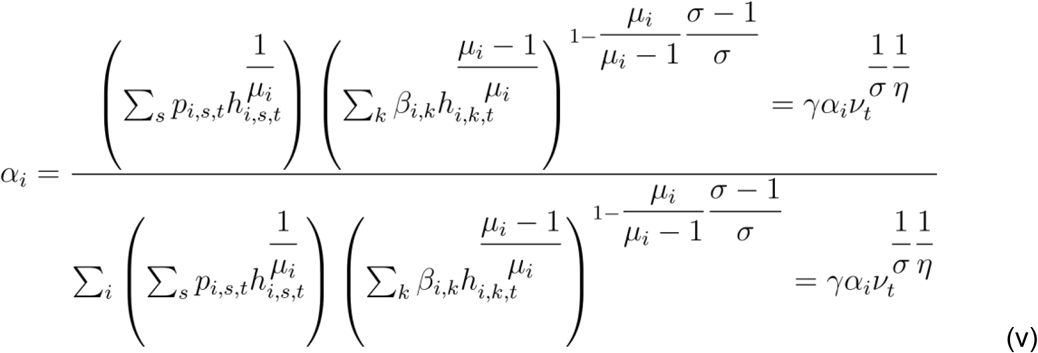

Note 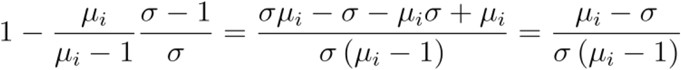

A2: Proof of Equation 11

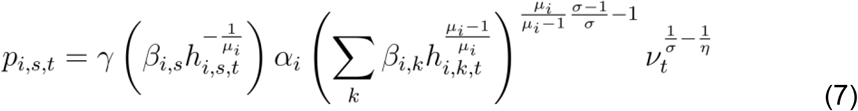

Using (6) we obtain:

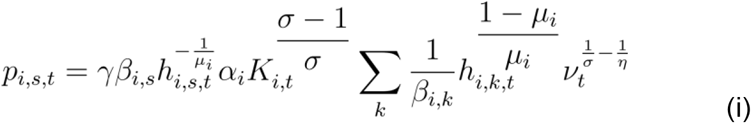

Multiplying (i) by *h*_*i,s,t*_,

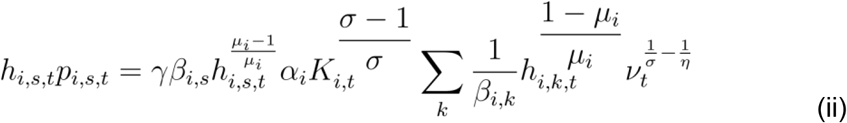

Summing over size classes and species,

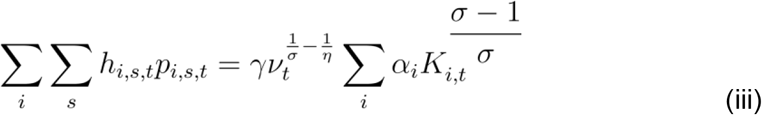

Using (5) we obtain,

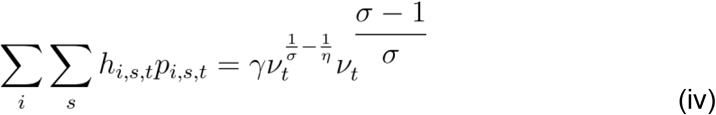

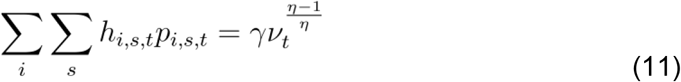

